# Dissecting AlphaFold’s Capabilities with Limited Sequence Information

**DOI:** 10.1101/2024.03.14.585076

**Authors:** Jannik Adrian Gut, Thomas Lemmin

## Abstract

Protein structure prediction, a fundamental challenge in computational biology, aims to predict a protein’s 3D structure from its amino acid sequence. This structure is pivotal for elucidating protein functions, interactions, and driving innovations in drug discovery and enzyme engineering. AlphaFold2, a powerful deep learning model, has revolutionized this field by leveraging phylogenetic information from multiple sequence alignments (MSAs) to achieve remarkable accuracy in protein structure prediction. However, a key question remains: how well does AlphaFold2 understand protein structures? This study investigates AlphaFold2’s capabilities when relying primarily on high-quality template structures, without the additional information provided by MSAs. By designing experiments that probe local and global structural understanding, we aimed to dissect its dependence on specific features and its ability to handle missing information. Our findings revealed AlphaFold2’s reliance on sterically valid C-*β* atoms for correctly interpreting structural templates. Additionally, we observed its remarkable ability to recover 3D structures from certain perturbations and the negligible impact of the previous structure in recycling. Collectively, these results support the hypothesis that AlphaFold2 has learned an accurate local biophysical energy function. However, this function seems most effective for local interactions. Our work significantly advances understanding of how deep learning models predict protein structures and provides valuable guidance for researchers aiming to overcome limitations in these models. protein folding, alphafold, side-chain, interpretability

## 1 Introduction

Proteins serve as nano-machines within cells, orchestrating a plethora of vital functions essential for life. Their remarkable versatility arises from their unique three-dimensional (3D) structures, dictated by the specific sequence of amino acids they are built from. This fundamental principle, known as Anfinsen’s dogma [3], underpins modern biology and fuels research in areas like drug discovery and enzyme engineering. However, progress in elucidating protein structures has been hindered by the labor-intensive nature of *in vitro* experiments required for atomic structure determination, resulting in only approximately 200,000 structures being resolved to date [7]. To surmount this bottleneck, researchers have increasingly turned to computational methodologies to unravel the intricacies of protein folding. Established in 1994, the Critical Assessment of Techniques for Protein Structure Prediction (CASP)[9] has played a crucial role in tracking advancements in this field. Recent years have been particularly transformative, fueled by two key factors: the exponential growth of sequential and structural protein data [8, 7] and the emergence of powerful machine learning methodologies, particularly deep learning, capable of harnessing this data more effectively. Notably, AlphaFold2 [14], a deep neural network unveiled in 2020, has revolutionized the field. This innovative model has achieved remarkable accuracy in predicting protein structures, marking a significant leap forward in our understanding of protein structure and function.

The AlphaFold2 pipeline follows a two-step process for predicting protein structures. First, it searches various protein sequence databases [24, 7, 8, 32, 23] using dedicated tools [31, 11, 17] to find similar sequences to the target protein across. This information gets compiled into a multiple sequence alignment (MSA), capturing the evolutionary relationships between the proteins. Simultaneously, AlphaFold2 identifies suitable 3D structures (templates) from closely related proteins to serve as initial structural models. These two sources of information, MSA and templates, are initially processed separately within the AlphaFold2 model. However, their representations are continuously refined through an iterative exchange of information, allowing the model to learn from both sources simultaneously. Finally, the refined representations of the MSA and templates are combined in the structure module of AlphaFold2 to generate the final protein structure and assign a confidence score (pLDDT) for each individual amino acid. The MSA has been observed to play a more significant role in predicting protein structure quality compared to templates. Notably, several pipelines leveraging AlphaFold2 [10, 35], and three out of the five default models of AlphaFold2, ignore the information from templates altogether. Intriguingly, contrary to previous pipelines, AF2Rank [27] showed that simply providing AlphaFold2 with a protein structure, without any sequence information, can be used to evaluate its plausibility and discriminate between real structures and decoys. The authors hypothesized that the structure module within AlphaFold2 has learned a robust biophysical energy function, and the MSA input might primarily serve to guide the model towards the correct energy minimum.

To further evaluate this hypothesis, we conducted an extensive investigation into the influence of the template input and structure recycling on AlphaFold2’s predictive accuracy. Through a comprehensive series of ablation studies, we assessed the model’s capability to reconstruct protein structures solely based on structure input, without relying on deep MSAs. Specifically, we examined AlphaFold2’s performance in side-chain packing and its resilience in recovering from artificially perturbed proteins. Our experimental code is openly accessible on GitHub^1^ and we have contributed new methods to the OpenFold [2] project^2^.

Our findings offer valuable insights for both AlphaFold2 users and developers who integrate the tool into their workflows. By enhancing the understanding of the model’s limitations and providing guidance on result interpretation, our work aims to empower users to critically assess results and leverage complementary tools when necessary. Additionally, it encourages exploration of existing tools or the development of innovative solutions to address these limitations, ultimately contributing to more accurate and reliable protein structure predictions.

## 2 Materials and methods

### 2.1 Datasets

The data used for the experiments was sourced from CASP13 [15] and CASP14 [16]. CASP13 was included in the analysis to evaluate potential bias and overfitting, given that AlphaFold2 was trained on proteins within the CASP13 dataset. Consequently, we anticipate observing higher scores compared to CASP14, which serves as a more realistic benchmark for assessing the model’s performance on unseen targets. This distinction is crucial for accurately gauging the model’s generalizability and effectiveness in predicting structures of novel protein sequences.

### 2.2 AlphaFold2

We primarily employed the LocalColabFold^3^ [21] implementation of AlphaFold2 in our experiments, as it offers a user-friendly command-line interface and faster inference enabled by the MMseqs webserver [22]. Given that AlphaFold2 largely ignores template information when provided with a deep MSA, a minimal MSA comprising only the query sequence (single sequence) was supplied to the model, unless otherwise specified. Furthermore, we developed OF2Rank, a protocol within the OpenFold [2] framework similar to AF2Rank that was previously used to assess the quality of protein structures [27]. This method involves replacing all original amino acid information in the template with glycine residues extended with a C-*β* atom. The sequence information is provided either as an all-gap multiple sequence alignment (*Gaps*) or as a single sequence MSA (*Single*). Due to technical constraints, OF2Rank was provided with the original amino acid sequence as an input sequence instead of gaps.

One key idea of AlphaFold2 is the recycling mechanism. This process iteratively refines protein structure predictions by feeding back the MSA embedding, pair embedding, and the structure prediction (*prev x*) of the previous iteration into the model. To evaluate the impact of a pre-existing template on predictions to this recycling, we modified OpenFold. This modification allows us to introduce a custom template structure as the “previous prediction” during the very first iteration of the recycling process (denoted as “-1”).

### 2.3 Side-chain packing

To assess AlphaFold2’s ability to reconstruct side-chains accurately, we designed four test cases using CASP13 and CASP14 data. In each case, all side-chain atoms were removed from the target proteins. In addition, to providing just the backbone as a template, we also positioned the C-*β* atom in three different ways:

i. *Non-informative C-β*: Placed next to the origin for a baseline comparison, ii. *Heuristic C-β*: Predicted using a rule-based approach based on the backbone atoms [27], iii. *Template C-β*: Maintained the original C-*β* position from the template

We extended our assessment to explore AlphaFold2’s ability to refine side-chain predictions using three external side-chain packing strategies: the lowest energy conformation provided by the widely used CHARMM 36 force field (*C36*), FASPR [12], a structure-based side-chain packing method and AttnPacker, a neural network-based side-chain packing method [19].

The modified templates and single sequences were then inputted into LocalColabFold without relaxation and without recycling, ensuring that the backbone remained more similar to the input template. Only the predictions from the first two AlphaFold2 models were considered, as they are the ones where template input is utilized.

### 2.4 Structural perturbation

Three distinct techniques were employed to generate controlled perturbations to the template structures (Figure 1). The refinement of perturbed structures was conducted using LocalColabFold with either a single sequence or MSA. In addition, we tested the newly implemented protocol within OpenFold to evaluate a more sequence-agnostic refinement.

**Figure 1:**
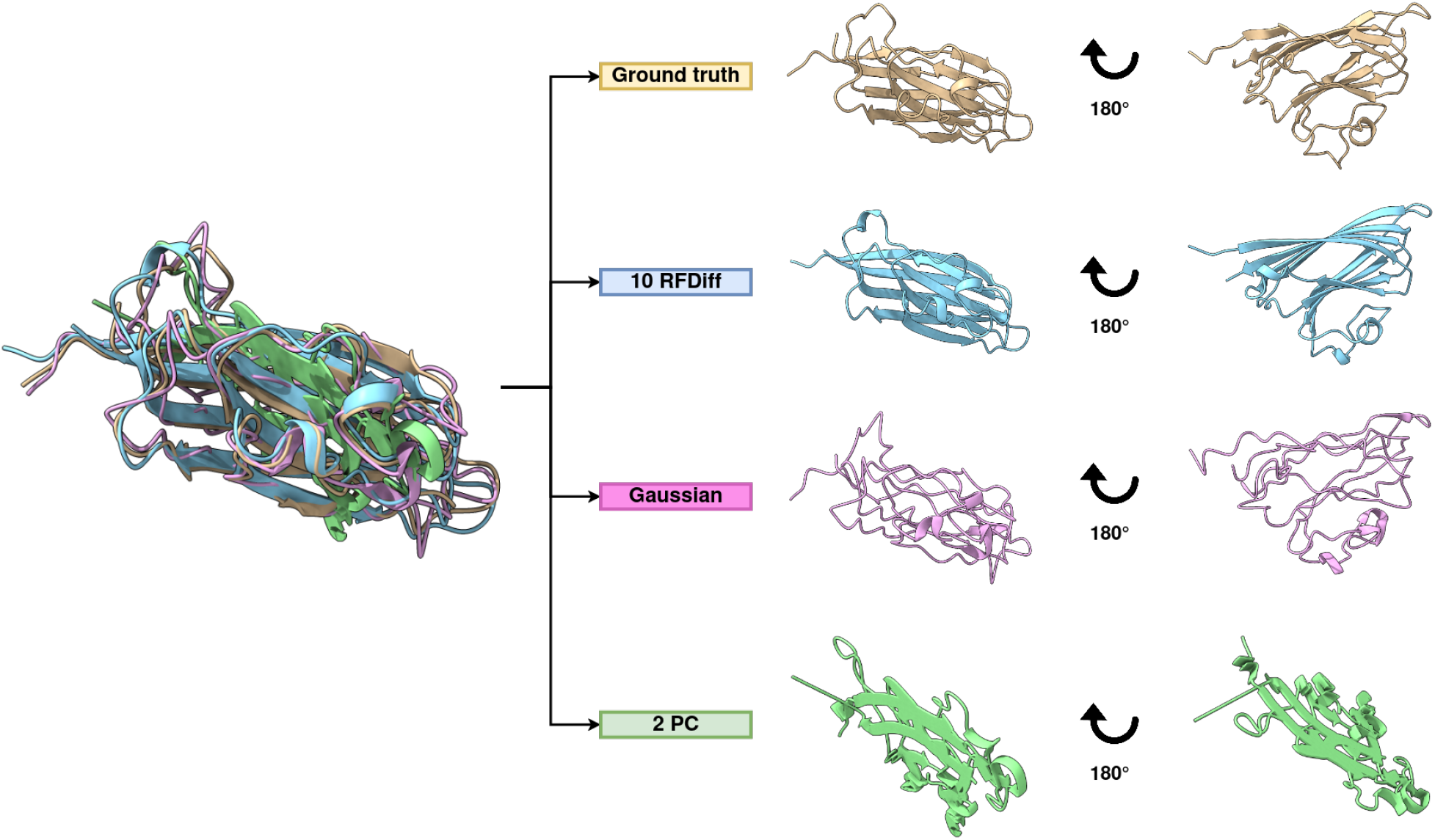
Illustration of structural perturbations. Ground truth protein structure (T1026, PDB id: 6s44) is shown in yellow, 10 partial RFdiffusion steps in blue, Gaussian noise in pink and 2D projection in green. All structures were rendered using ChimeraX [20].

#### 2.4.1 Gaussian noise

The simplest perturbation method involved adding Gaussian noise to the template coordinates. We independently sampled values from a standard normal distribution (mean = 0Å) for each atomic coordinate dimension. A predefined standard deviation of 1Å determined the magnitude of the introduced noise. These sampled values were then added to the original coordinates, introducing controlled deviations from the initial structure.

#### 2.4.2 Principal component analysis

Principal Component Analysis (PCA) [6] was employed to transform and project the protein into an orthogonal reference frame that explains the most variance. Separate PCAs were computed for each protein in the CASP13 and CASP14 datasets. Subsequently, each structure was projected onto the sub-space defined by the first principal component (*1 PC*) or the first two principal components (*2 PC*). This effectively captured the major structural variations within each protein structure.

#### 2.4.3 RFdiffusion

RFdiffusion [36] leverages diffusion techniques [30] to generate new proteins from noise. Instead of executing the complete diffusion process, which generates entirely new backbones conformations, we ran RFdiffusion for a limited number of partial steps (1, 5 and 10 steps) [33]. This strategy produced backbones that remained closer to the starting structure, with the degree of variability increasing with the number of steps. Subsequently, side-chains were reconstructed using FASPR or AttnPacker.

### 2.5 Evaluation metrics

To assess the accuracy of the predicted protein structures, we employed a variety of metrics. These metrics can be broadly categorized into three classes based on the level of detail they capture.

The first class of accuracy metrics focuses solely on the *α*-carbon atom from the backbone. This includes the Template Modeling (TM)-score [37, 5], which quantifies the structural similarity between predicted and reference structures, and the *α*-RMSD [5], measuring the difference between the C-*α* positions of the predicted and reference structures after optimal superposition using the Kabsch algorithm.

The next class of accuracy metrics encompasses RMSD and the Local Distance Difference Test (lDDT) [18, 5], which evaluates all atom pairs (backbone and side-chain) within a predefined radius excluding those belonging to the same residue. In addition, AlphaFold2 predicts a score called pLDDT, which estimates the lDDT value of its generated structures and helps guide the selection of the best model among multiple predictions. Individual perresidue lDDT scores can be combined to generate a single, global lDDT score. To accommodate potentially invalid protein structures with steric clashes, stereochemical checks were disabled during lDDT calculations.

The last class of metrics evaluates the accuracy of side-chain packing with the Mean Absolute Error (MAE) for the first four dihedral angles of the side-chains [19].

Pdb-tools [26], the MAXIT suite, ProDy [4], Scikit-learn [25] were used for implementing the experiments and analyzing the results.

## 3 Results

To assess AlphaFold2’s [14] understanding of protein structure, we designed two complementary tasks. The first task assessed AlphaFold2’s local understanding by testing its ability to rebuild and pack side-chains onto a provided backbone template. This evaluated AlphaFold2’s ability to handle individual amino acid interactions within the protein structure. The second task investigated the global understanding by evaluating AlphaFold2’s effectiveness in recovering the correct protein structure from a perturbed template. Previous studies have indicated that AlphaFold2 largely disregards template input when provided with a deep multiple sequence alignment (MSA) [1]. Therefore, to isolate its capability in utilizing structural information, we primarily employed a single sequence along with the template as input for AlphaFold2 in most experiments. To assess potential overfitting concerns, we compared results on proteins from CASP13 (part of AlphaFold2’s training data) to those from CASP14 (unseen data). Since we did not observe any major differences between these two datasets, we will focus on presenting the results from CASP14 in this section, but report data of both datasets in supplementary information Section A and supplementary information Section C for side-chain packing and structural refinement respectively.

### 3.1 Side-chain packing

Side-chain packing is a critical process in protein structure prediction, involving the prediction of the three-dimensional positions of amino acid side-chains relative to the protein backbone. This task is essential for accurate modeling of protein structures and for understanding their biological functions.

Our initial evaluation focused on AlphaFold2’s ability to pack side-chains using only the backbone atoms and different approaches for the placement of C-*β* atoms: all close to the origin (*Non-informative C-β*), using a heuristic (*Heuristic C-β*), or preserving the correct position from the template (*Template C-β*). When the template lacked C-*β* information (either missing or Non-informative C-*β*), the predicted structures suffered significant accuracy loss. The average TM-score dropped to approximately 0.41 ± 0.20 and 0.32 ± 0.20, respectively, indicating a failure to preserve the 3D structure. Providing more informative C-*β* positions, either through heuristics or the template, significantly improved the results. The average TM-score remained very high (nearly 0.97 ± 0.03) when using the heuristic C-*β* placement, indicating minimal alterations of the backbone structure from the template. It’s worth noting that this score is higher than the average TM-score achieved by AlphaFold2 predictions using a full MSA without a template (approximately 0.8 ± 0.18).

These high TM-scores suggest that side-chain packing metrics can be reliably analyzed, since the overall protein fold is mostly maintained. The heuristic C-*β* placement achieved a promising average lDDT score of 0.89 ± 0.05, exceeding the baseline method (standard pipeline with full MSA) which had an average lDDT of 0.74 ± 0.15. Next, we assessed AlphaFold2’s potential to enhance the predictions of three different methods for side-chain placement: predefined conformations from the CHARMM36 force field, FASPR [12], and AttnPacker [19]. We employed the AutoPSF plug-in from VMD [13] to assign a single, predefined side-chain structure to each residue based on the CHARMM36 force field (*C36*). While fast and simple, it neglects the local environment, leading to clashes and sub-optimal packing. FASPR [12] utilizes a tree search algorithm to identify energetically favorable placements for predefined rotamers [28]. This approach offers better balance between speed and accuracy by using a predefined library of conformations but allowing for optimization based on energy minimization. Finally, AttnPacker is a deep learning-based method that directly predicts side-chain coordinates without relying on predefined rotamers. This approach offers greater flexibility in side-chain placement compared to rotamer-based methods.

Refining the repacked protein structures through AlphaFold2 resulted in a slight decrease in the average TM-score to 0.98 ± 0.03, indicating again minimal backbone alteration. Interestingly, refining the packing with AlphaFold2 significantly boosted the average lDDT scores when the initial packing was poor (e.g., predefined rotamers from the CHARMM36 force field). However, for methods like FASPR and AttnPacker, which already produced good initial packing, the lDDT scores remained similar or even decreased slightly after AlphaFold2 refinement (Figure 2).

**Figure 2:**
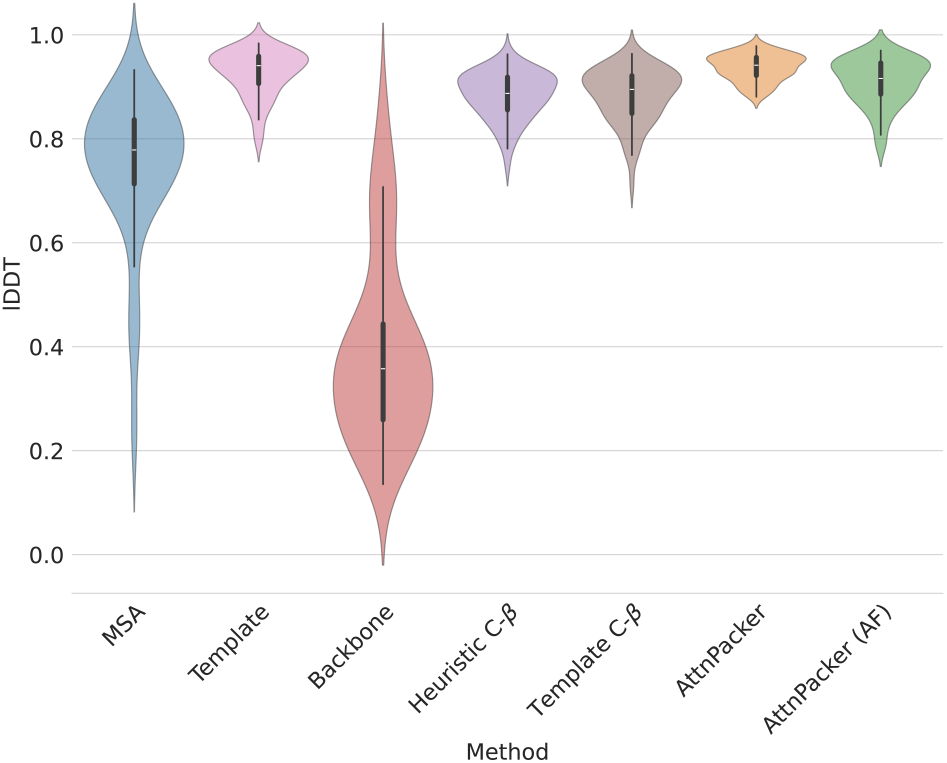
Average lDDT scores for side-chain packing task. Violin plots show the distribution of average lDDT scores for various AlphaFold2 template configurations. *MSA*: full MSA, but no template, *Template*: full template, *Backbone*: the protein backbone *Heuristic C-β*: template with heuristically placed C-*β, Template with C-β*: backbone and C-*β* from the ground truth, *AttnPacker* and *AttnPacker (AF)*: ground truth backbone with side-chains placed by AttnPacker before and after AF refinement

Analyzing the mean RMSD revealed similar effects of AlphaFold2 refinement on different packing methods. Notably, competitive side-chain packing methods like FASPR and AttnPacker see a marginal change in performance after post-processing with AlphaFold2. Conversely, when using only the default low energy conformation from the CHARMM36 force field, which initially performs poorly (RMSD: 1.77 Å ± 0.15), exhibited a significant improvement in side-chain placement (RMSD: 0.88 Å ± 0.25) after AlphaFold2 refinement. When provided with only the heuristic C-*β*, AlphaFold2 is able to predict the side-chain position with a similar precision as FASPR, RMSD 0.91 Å ± 0.25 and 0.89 Å ± 0.27 respectively. While providing the correct C-*β* information leads to a marginal improvement (RMSD: 0.86 Å ± 0.27) over the heuristic approach.

Furthermore, we noticed a slight decrease in confidence and precision as residues become more exposed at the protein surface (Section B in the supplementary information). This decrease is consistent across all template-informed protocols, but is notably steeper for the standard AlphaFold2 protocol relying solely on MSAs.

### 3.2 Structural refinement

Next, we assessed AlphaFold2’s ability to refine perturbed templates. Three perturbation protocols were implemented and tested: introducing random Gaussian noise to atom coordinates (*Gaussian*), projecting the entire structure onto a 1D or 2D space defined by principal components (*1 PC* and *2 PC* respectively), and partially denoising the structure using RFdiffusion (*RFDiff*). The full table with scores can be found in supplementary information Section C.

Despite struggling to fully recover the original structure for several perturbations, AlphaFold2 generated sterically valid structures in most cases. This is evident in the necessity to disable stereochemical checks from OpenStructure [5] when calculating lDDT for templates perturbed with the Gaussian noise and projected in to the PCA eigenspace. These checks ensure proper atomic interactions, and the initial violation by a majority of residues suggests significant structural distortion. However, the fact that these checks would not be needed for post-processed structures implies that AlphaFold2, while not achieving complete fold recovery, produces structures with proper atomic interactions.

AlphaFold2 demonstrated good recovery capabilities for templates perturbed with Gaussian noise (Figure 3). The average TM-score increased from 0.90 ± 0.05 to 0.92 ± 0.06, and the lDDT score improved from 0.66 ± 0.00 to 0.82 ± 0.07. Notably, AlphaFold2 excelled at recovering structures projected onto the two-dimensional PCA space. Here, refinement significantly boosted accuracy, with the average TM-score rising from a low 0.49 ± 0.08 to 0.86 ± 0.15 and the mean lDDT increasing from 0.53 ± 0.09 to 0.79 ± 0.12. While refinement also improved results for one-dimensional projections, the final refined scores (average TM-score: 0.44 ± 0.21 and mean lDDT: 0.39 ± 0.20) remained relatively low.

**Figure 3:**
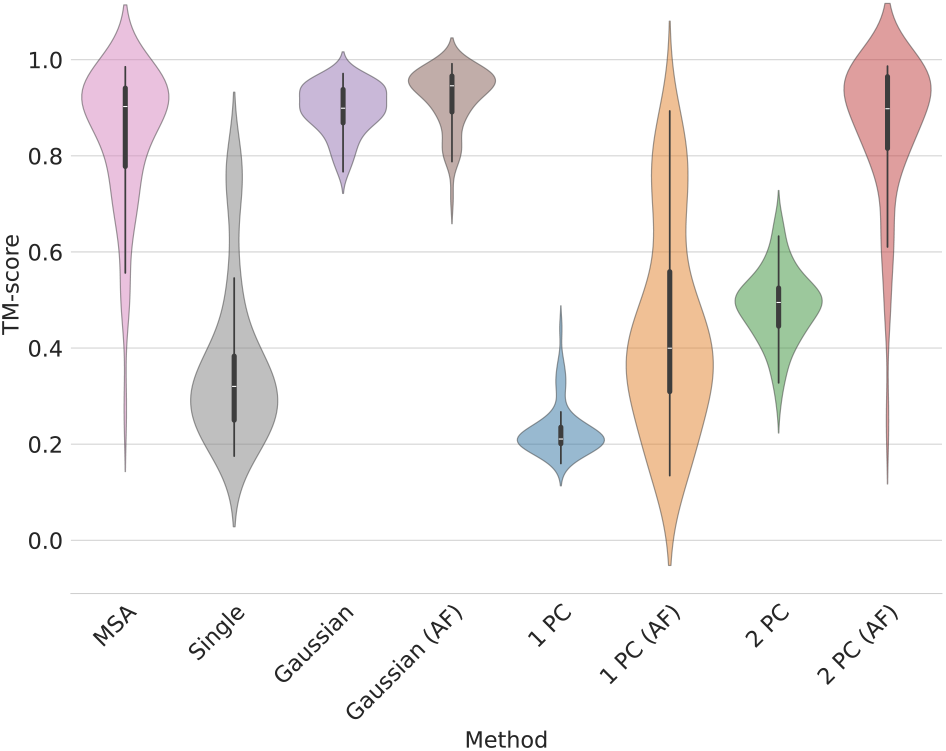
Violin plot of selected perturbation strategies. *(AF)* is used to identify the post-processing by AlphaFold2, *MSA* uses the vanilla AlphaFold2 pipeline with a full MSA and no pipelines, while *Single* uses only the single sequence, *Gaussian* adds Gaussian noise to the coordinates, *1 PC* reduces proteins to the first principal component, *2 PC* reduces targets to the first two principal components

For the RFdiffusion based perturbation, we observed a gradual decrease in both TM-score and lDDT, as the number of partial diffusion steps increased. The average TM-score dropped from 0.97 ± 0.01 with one diffusion step to 0.87 ± 0.05 with 10 steps (Figure 4), while the average lDDT fell from 0.80 ± 0.02 to 0.67 ± 0.04. These scores suggest that despite the perturbation, the protein remained within the same general fold. This implies that the starting point for refinement by AlphaFold2 was still favorable for recovery of the correct structure, rather than collapsing into an alternative fold. Interestingly, while the average TM-score remained relatively unchanged after AlphaFold2 refinement for all diffusion steps, the average lDDT score consistently improved by around 0.04. This finding suggests that in this configuration, AlphaFold2 primarily refines local structures, with minimal adjustments to the protein backbone, as reflected by the stable TM-scores.

**Figure 4:**
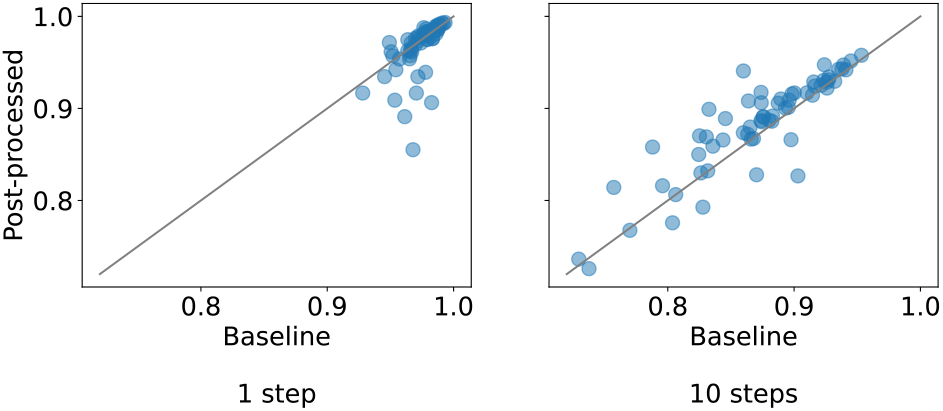
Change in TM-score after AlphaFold2 refinement of RFdiffusion-perturbed templates. Each point represents a single target from the CASP14 dataset. The TM-score of the RFdiffusion-perturbed structure before refinement is displayed on the x-axis, with separate plots for 1 and 10 partial steps on the left and right, respectively. The y-axis shows the TM-score after refinement using AlphaFold2. The side-chains were packed using FASPR.

Previous research has shown that AlphaFold2 can estimate template quality more precisely by replacing all residues in the template with glycine extended with a C-*β* atom, and providing the sequence with all gaps and an empty multiple sequence alignment (AF2Rank) [27]. We implemented a similar method within OpenFold, which we term OF2Rank. Interestingly, while OF2Rank achieved lower performance compared to AlphaFold when recovering structures from templates perturbed with Gaussian noise or projected into 1D/2D PCA space, it showed comparable effectiveness for RFdiffusion-perturbed templates. Furthermore, the advantages of OF2Rank became more pronounced with increasing numbers of RFdiffusion steps. At 10 partial diffusion steps, OF2Rank increased the average lDDT reached by 0.03. Intriguingly, using an all-gap MSA yielded slightly better predictions compared to the single sequence MSA, mirroring observations from AF2Rank [27]. The average difference between the all-gap and single sequence pipelines for both average TM-score and average lDDT was approximately 0.03 each. However, overall, AlphaFold pipeline generally outperformed OF2Rank for the refinement of perturbed templates.

Replacing the template input with prev x had minimal impact on the predictions, with the TM-Score increasing from 0.38 to 0.43 (Appendix D). Further tests with full MSAs and disabled prev x showed no significant difference (TM-Score of 0.84 ± 0.15 for both with and without prev x in CASP14).

## 4 Discussion

This study investigated capabilities and limitations of AlphaFold2’s ability to understand protein structures. We designed a series of experiments to investigate how AlphaFold2 handles both local features, like side-chain packing, and global features, like perturbations to the backbone.

Our findings suggest that C-*β* atoms are crucial for AlphaFold2 to recognize a template as a valid protein structure. When C-*β*s are present, AlphaFold2 prioritizes side-chain packing and only marginally alters the backbone if the phylogenic signal is weak. This could be leveraged as pre-processing pipeline that refines incomplete experimental structures by standardizing atom numbering and modeling missing residues. Interestingly, a simple heuristic for C-*β* placement achieves performance comparable to providing the original position of the C-*β*. However, providing more side-chain information of seemingly lower quality, like the predefined conformations from the CHARMM36 force field, did not significantly improve the side-chain packing performance. Furthermore, pre-packing the template with dedicated side-chain packing algorithms like FASPR and AttnPacker only marginally impacted the final packing performance by AlphaFold2. This would further suggest that AlphaFold2’s understanding of the protein structure relies more on the presence of stereochemically valid C-*β* atoms than the detailed packing of the entire side-chain.

We also observed that AlphaFold2’s side-chain packing performance remained nearly consistent regardless of residue burial depth, while the performance dropped significantly for surface residues when AlphaFold2 relied solely on the MSA. This finding underlines the importance of high-quality templates, especially when dealing with shallow MSAs.

Perturbing the input template with various methods revealed insightful details regarding AlphaFold2’s capabilities in recovering three-dimensional protein structures. AlphaFold2 efficiently recovered structures perturbed with Gaussian noise, which primarily involves local adjustments to bond lengths and angles within residues. This ease of recovery hints at AlphaFold2 potentially having learnt a biophysical energy model for proper steric interactions, similar to classical optimization methods that utilize molecular force fields. Even more striking is AlphaFold2’s ability to recover details of a three-dimensional protein structure from 2D-like templates. This suggests that AlphaFold2 can effectively navigate the transition from a limited structural representation to a full 3D structure. Research on OpenFold, a reimplementation of AlphaFold2, showed that it first predicts a 2D representation during early stages of training before transitioning to 3D [2]. This finding raises the intriguing possibility that AlphaFold2 has preserved a similar internal representation of a protein structure.

A significant difference in relative performance was observed between AF and the OF2Rank method when refining templates perturbed with Gaussian noise or projected into 1D/2D PCA space, compared to those perturbed with RFdiffusion. The key distinction lies in the nature of the perturbations. RFdiffusion typically introduces realistic modifications that maintain valid protein structures, whereas other methods often generate structures with steric clashes and other imperfections.

The markedly lower performance of OF2Rank on these unrealistic protein structures highlights a potential strength of the underlying approach. By struggling with these templates, OF2Rank might be more adept at identifying unreliable starting points. This aligns with the notion that AlphaFold2 was trained on the assumption of structurally sound templates. When presented with corrupted starting structures and lacking a reliable MSA for guidance, AlphaFold2’s performance will decline as it grapples with reconciling the conflicting information.

Our observations of minimal impact from structure recycling in AlphaFold2 align with the decision by AlphaFold3’s authors to remove this mechanism entirely [**?**].

Prior research on AlphaFold2 using MSA-free protocols suggests the structure module might have learned a valid biophysical energy function, independent of MSAs [27]. AlphaFold2 would then act as an unrolled optimizer and make iterative adjustments guided by the learned potential to find a low-energy state, corresponding to a refined protein structure. Our results support this hypothesis, but suggest limitations. This optimization likely operates within a restricted neighborhood that requires both the backbone and stereochemically valid C*-β* atoms. Additionally, the function appears most effective for local molecular interactions, as evidenced by its successful handling of side-chain packing and Gaussian noise perturbations.

Our work provides valuable guidance for users to critically evaluate AlphaFold2’s predictions and identify scenarios where complementary tools might be necessary. These insights pave the way for further exploration of existing methods or the development of novel strategies to overcome AlphaFold2’s limitations, ultimately leading to more robust and reliable protein structure prediction.

## 6 Author contributions statement

J.A.G. and T.L. conceived the experiment(s), J.A.G. conducted the experiment(s), J.A.G. and T.L. analyzed the results. J.A.G. and T.L. wrote and reviewed the manuscript.

## 7 Acknowledgements

We thank Axel Giottonini for feedback on the manuscript.

This work is supported by funds from the FreeNovation 2023 grant and the Swiss National Science Foundation (PCEFP3 194606).

## A Side-chain packing table

Table A1 shows the results of the different side-chain packing experiments. The scores are computed for each target independently, then average and standard deviation are determined over the target scores. The major results are discussed in Section 3.1 of the main text.

**Figure A1:**
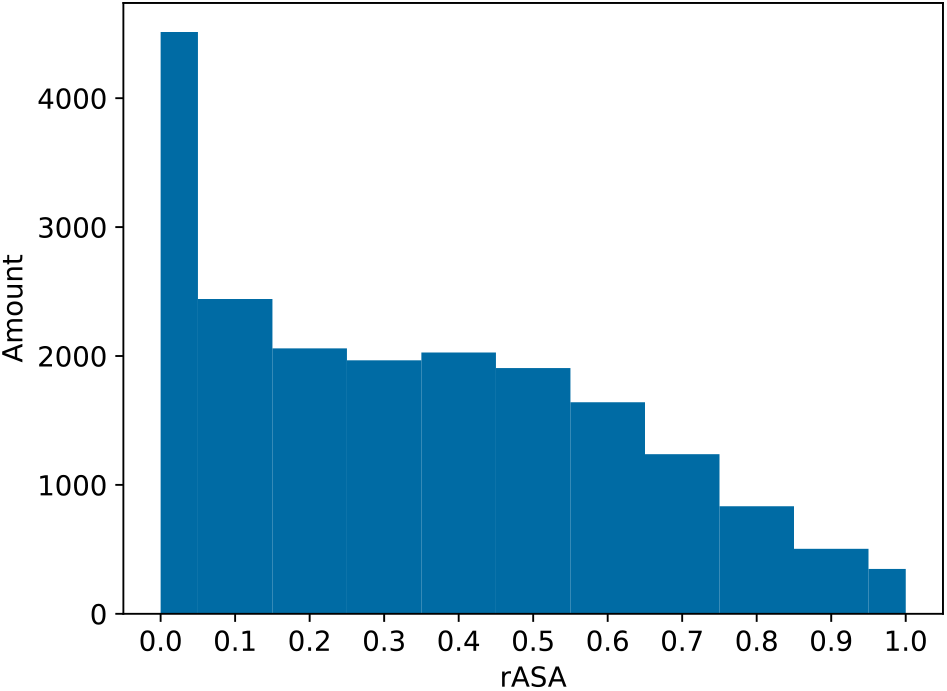
Histogram of rASA bins on the CASP13 dataset.

## B Side-chain packing performance correlation with other evaluation measures

To further analyze the results of the side-chain packing experiment, a comparison with confidence (pLDDT) and their relative accessible surface area (rASA)[29, 34] is performed. The residues are binned into 11 bins depending on the rASA rounded to the first decimal in the ground truth target. Therefore, the histogram for CASP13, depicted in Figure A1, and CASP14, shown in Figure A2, is the same for each side-chain packing method with the same dataset. A plot comparing rASA with pLDDT and lDDT on the CASP13 dataset can be seen in Figure A3 and a similar plot for CASP14 can be found in Figure A4.

**Figure A2:**
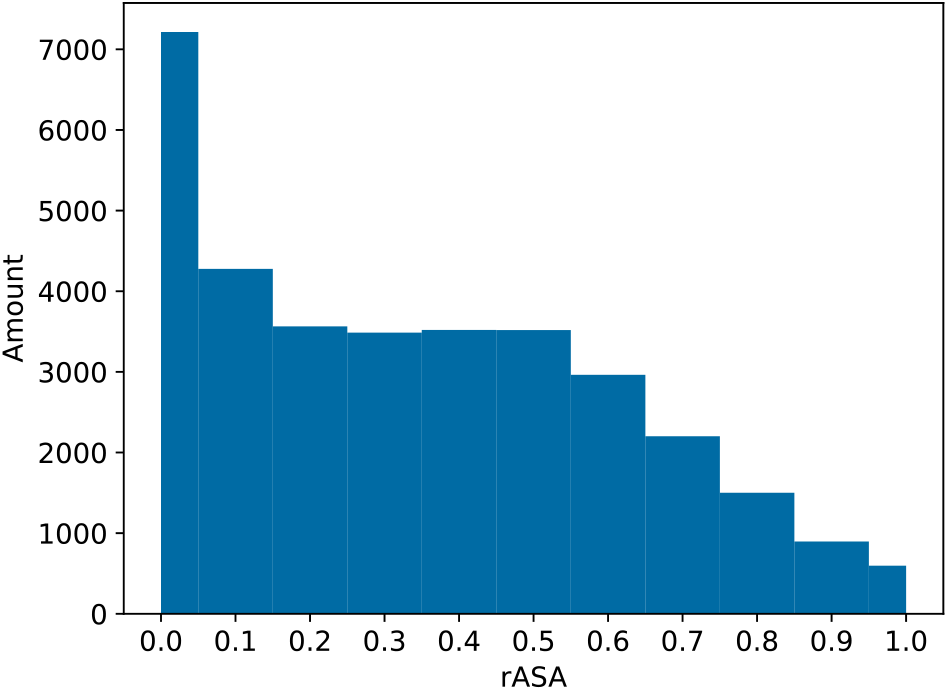
Histogram of rASA bins on the CASP14 dataset.

As expected, the pLDDT and lDDT drops with increasing rASA on average. This drop is only minor until the bins at around a rASA of 0.9, where the drop becomes steeper. Comparing the different packers, the majority of them stay pretty well together; their scores are ordered the same as in the results in Section 3.1. The lDDT and pLDDT have very similar curves, which indicates that pLDDT is also a good lDDT estimator in these circumstances. The exact Pearson correlation coefficients are reported in Table A2 between lDDT, pLDDT and rASA on the CASP13 and the CASP14 dataset for multiple side-chain placement protocols.

The first outlier to the majority is AttnPacker, which generally has a lower confidence, but a higher lDDT. The reason for the lower score in the pLDDT figure is due that AttnPacker reports its own confidence, which is not the same as the confidence from AlphaFold2. The higher lDDT score then can be explained by a superior packing performance and that the backbone, which influences this metric, did not get modified.

The other outlier is vanilla AlphaFold2 using the full multiple sequence alignment (MSA). This setup performs worse than the backbone informed methods, but the pLDDT and lDDT still have a good correlation.

**Figure A3:**
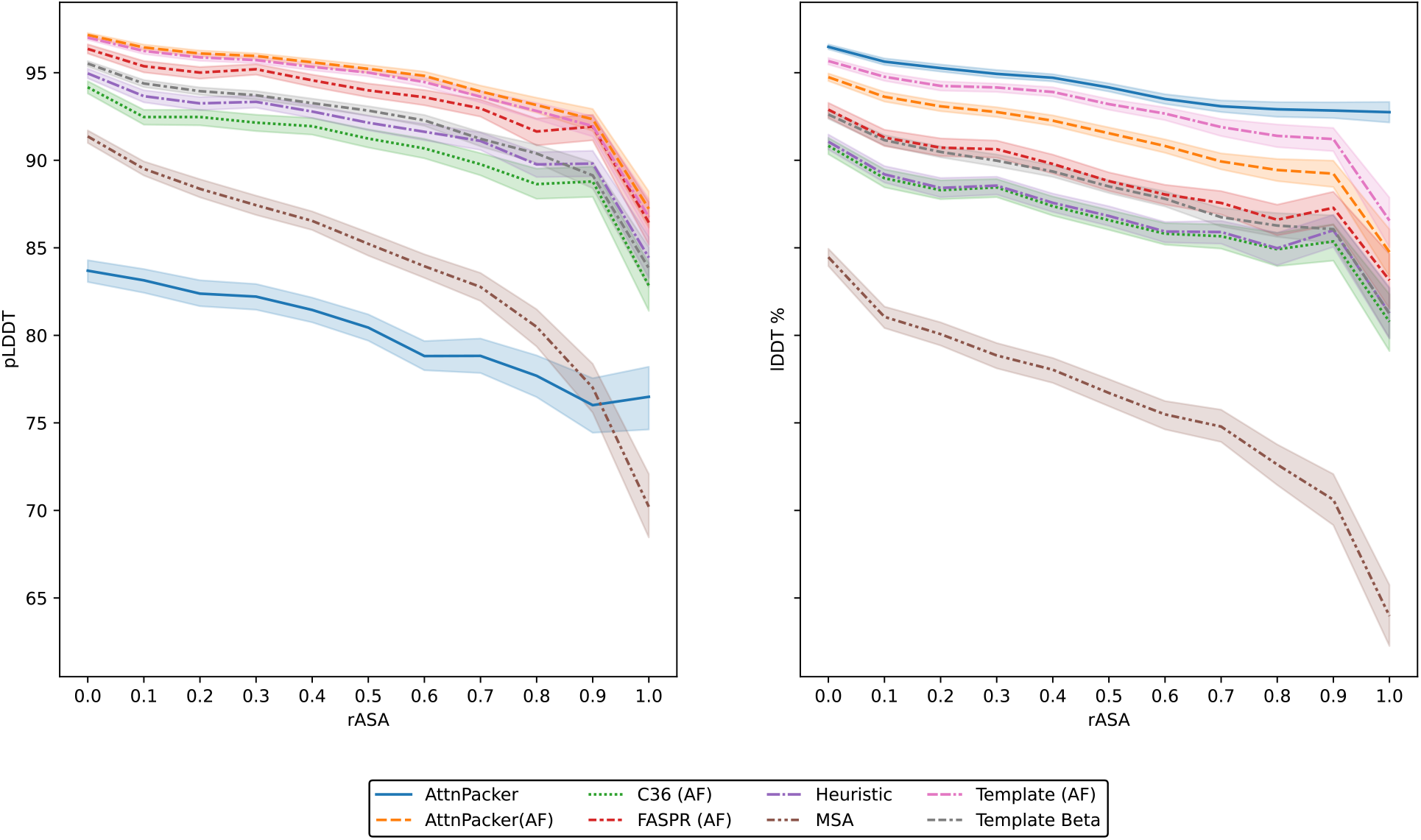
Relationship between rASA and pLDDT and lDDT for the CASP13 dataset. The line indicates the average score and the shadow area shows the 95% confidence interval. Side-chain packing using: ground truth backbone and side-chains placed with AttnPacker (*AttnPacker*), CHARMM 36 force field (*C36*) or FASPR (*FASPR*), ground truth backbone and the C-*β* placed with a heuristic (*Heuristic*), AlphaFold2 with a full MSA and no template (*MSA*), full ground truth template (*Template (AF)*), backbone and C-*β* from ground truth template (*Template Beta*). The *(AF)* suffix indicates AlphaFold2 post-processing after side-chain packing.

**Figure A4:**
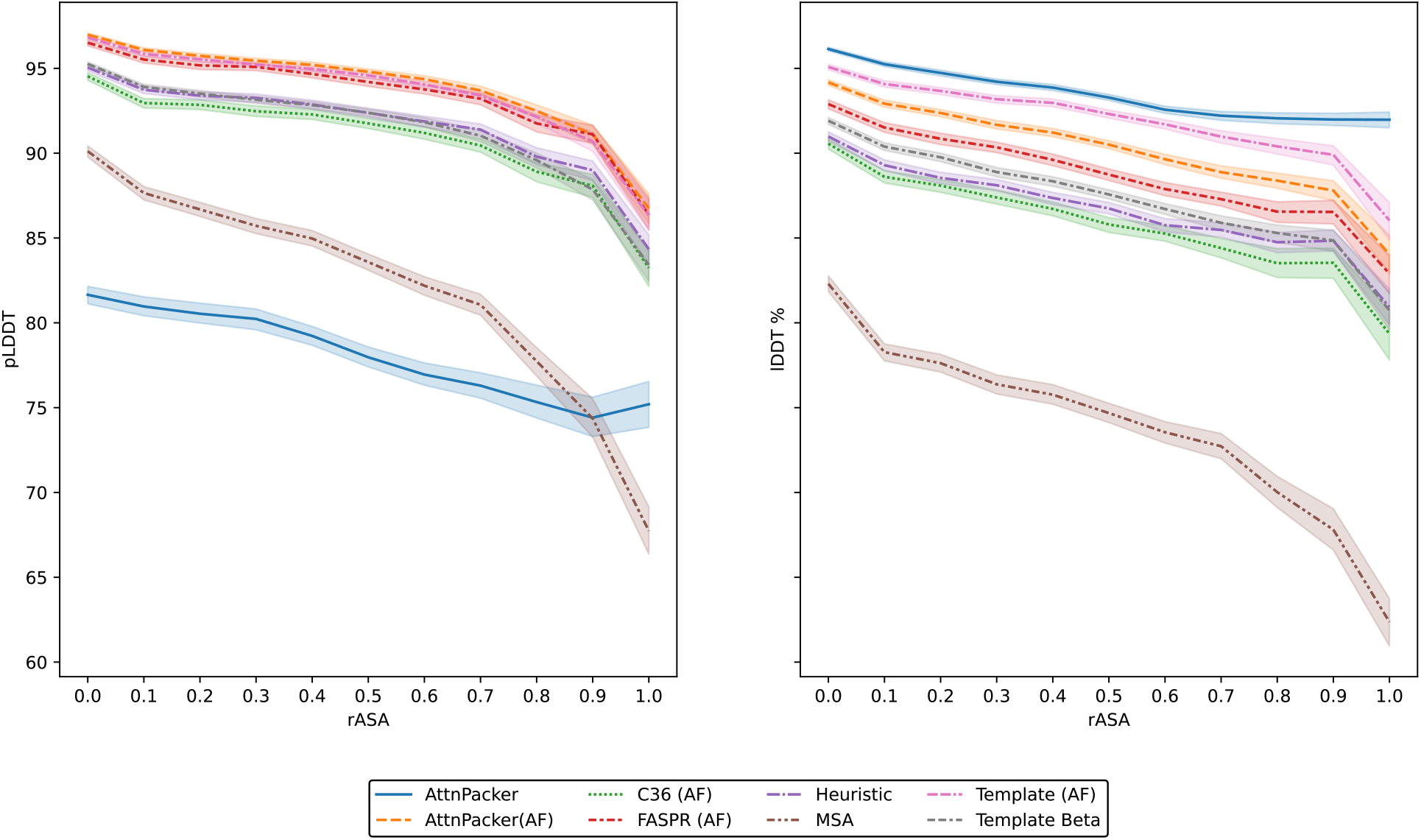
Relationship between rASA and pLDDT and lDDT for the CASP14 dataset.The line indicates the average score and the shadow area shows the 95% confidence interval. Side-chain packing using: ground truth backbone and side-chains placed with AttnPacker (*AttnPacker*), CHARMM 36 force field (*C36*) or FASPR (*FASPR*), ground truth backbone and the C-*β* placed with a heuristic (*Heuristic*), AlphaFold2 with a full MSA and no template (*MSA*), full ground truth template (*Template (AF)*), backbone and C-*β* from ground truth template (*Template Beta*). The *(AF)* suffix indicates AlphaFold2 post-processing after side-chain packing.

**Table A1:** Side-chain packing results.

| Method | Dataset | TM-score ↑ | IDDT ↑ | RMSD ($\text{\AA}$) ↓ | MAE 1 (rad) ↓ | MAE 2 (rad) ↓ | MAE 3 (rad) ↓ | MAE 4 (rad) ↓ |
| --- | --- | --- | --- | --- | --- | --- | --- | --- |
| **MSA** | **CASP13** | 0.833±0.195 | 0.812±0.113 | 0.797±0.224 | 0.516±0.187 | 0.499±0.130 | 0.817±0.162 | 0.947±0.233 |
| | **CASP14** | 0.793±0.178 | 0.743±0.146 | 0.973±0.254 | 0.632±0.191 | 0.573±0.149 | 0.882±0.158 | 0.877±0.229 |
| **Template** | **CASP13** | 0.985±0.032 | 0.945±0.038 | 0.498±0.200 | 0.273±0.149 | 0.315±0.128 | 0.586±0.180 | 0.795±0.229 |
| | **CASP14** | 0.982±0.031 | 0.926±0.045 | 0.616±0.235 | 0.350±0.185 | 0.381±0.162 | 0.649±0.181 | 0.756±0.235 |
| **Backbone** | **CASP13** | 0.412±0.204 | 0.393±0.166 | 1.107±0.170 | 0.739±0.139 | 0.629±0.105 | 0.910±0.143 | 0.957±0.212 |
| | **CASP14** | 0.409±0.200 | 0.392±0.178 | 1.183±0.198 | 0.784±0.143 | 0.653±0.107 | 0.932±0.152 | 0.875±0.234 |
| **Non-informative C-β** | **CASP13** | 0.323±0.200 | 0.304±0.174 | 1.164±0.167 | 0.785±0.141 | 0.653±0.102 | 0.916±0.165 | 0.953±0.221 |
| | **CASP14** | 0.318±0.210 | 0.311±0.189 | 1.238±0.184 | 0.828±0.137 | 0.673±0.098 | 0.950±0.163 | 0.876±0.238 |
| **Heuristic C-β** | **CASP13** | 0.970±0.093 | 0.895±0.070 | 0.785±0.209 | 0.489±0.173 | 0.500±0.123 | 0.819±0.158 | 0.939±0.230 |
| | **CASP14** | 0.979±0.033 | 0.883±0.045 | 0.910±0.247 | 0.572±0.190 | 0.557±0.156 | 0.870±0.170 | 0.869±0.232 |
| **Template C-β** | **CASP13** | 0.974±0.062 | 0.905±0.040 | 0.735±0.218 | 0.460±0.176 | 0.464±0.129 | 0.794±0.169 | 0.935±0.221 |
| | **CASP14** | 0.971±0.038 | 0.883±0.052 | 0.860±0.265 | 0.544±0.196 | 0.516±0.164 | 0.842±0.165 | 0.854±0.241 |
| **C36** | **CASP13** | 1.000±0.000 | 0.782±0.017 | 1.731±0.149 | 1.339±0.086 | 0.732±0.100 | 0.997±0.187 | 0.951±0.213 |
| | **CASP14** | 1.000±0.000 | 0.565±0.076 | 1.769±0.154 | 1.315±0.096 | 0.743±0.121 | 0.990±0.168 | 0.869±0.240 |
| **C36 (AF)** | **CASP13** | 0.962±0.111 | 0.890±0.099 | 0.778±0.215 | 0.485±0.177 | 0.475±0.127 | 0.817±0.191 | 0.948±0.222 |
| | **CASP14** | 0.978±0.036 | 0.873±0.057 | 0.883±0.251 | 0.551±0.190 | 0.527±0.153 | 0.849±0.158 | 0.860±0.242 |
| **FASPR** | **CASP13** | 1.000±0.000 | 0.926±0.027 | 0.756±0.228 | 0.451±0.167 | 0.488±0.140 | 0.863±0.180 | 1.017±0.279 |
| | **CASP14** | 1.000±0.000 | 0.911±0.030 | 0.896±0.265 | 0.548±0.197 | 0.554±0.168 | 0.860±0.156 | 0.963±0.246 |
| **FASPR (AF)** | **CASP13** | 0.975±0.088 | 0.918±0.069 | 0.697±0.234 | 0.420±0.177 | 0.451±0.140 | 0.802±0.194 | 0.931±0.238 |
| | **CASP14** | 0.983±0.030 | 0.905±0.043 | 0.834±0.274 | 0.511±0.206 | 0.514±0.176 | 0.820±0.152 | 0.888±0.237 |
| **AttnPacker** | **CASP13** | 1.000±0.000 | 0.953±0.025 | 0.531±0.201 | 0.301±0.149 | 0.393±0.126 | 0.775±0.188 | 0.951±0.225 |
| | **CASP14** | 1.000±0.000 | 0.937±0.026 | 0.687±0.220 | 0.414±0.176 | 0.480±0.151 | 0.832±0.159 | 0.891±0.229 |
| **AttnPacker (AF)** | **CASP13** | 0.986±0.031 | 0.933±0.038 | 0.614±0.220 | 0.355±0.175 | 0.405±0.131 | 0.733±0.184 | 0.918±0.226 |
| | **CASP14** | 0.983±0.030 | 0.910±0.045 | 0.772±0.263 | 0.461±0.203 | 0.484±0.163 | 0.790±0.164 | 0.862±0.235 |

Results of the side-chain packing experiment. Averages with standard deviation are shown for CASP13 and CASP14 separately. TM-score is used to score the backbone, while IDDT and RMSD score backbone and side-chains simultaneously and the mean absolute errors of dihedral side-chain angles in Radians, starting from the first angle to the fourth give exclusive side-chain results. AlphaFold2 refinement with: a full MSA and no template (*MSA*), full ground truth template (*Template*), just the backbone (*Backbone*), ground truth backbone and C-β placed next to the origin *Non-informative C- $\beta$* , ground truth backbone and the C-β placed with a heuristic (*Heuristic C- $\beta$* ) and backbone and C-β from ground truth template (*Template C- $\beta$* ), or ground truth backbone and side-chains placed with CHARMM 36 force field (*C36*), FASPR (*FASPR*) or AttnPacker (*AttnPacker*). The (*AF*) suffix indicates AlphaFold post-processing after side-chain packing.

## C Refinement tables

The major results of the refinement task are discussed in Section 3.2 in the main text. Table A3 shows results for Gaussian noise and principal component reduction, while Table A4 displays the results of RFdiffusion. The scores are computed for each target independently, then average and standard deviation are determined over the target scores.

## D Prev x experiments

Since prev x requires complete structures, all PDB entries with missing residues were excluded from the analysis presented in Table A5. This filtering resulted in a dataset of 60 structures for CASP13 and 48 structures for CASP14. For comparability, the scores for the other experiments were recomputed for this subset.

These experiments were conducted using a custom build of OpenFold. This version utilizes pre-trained weights and offers the ability to disable the prev x output or provide a structure for the recycling input during the first pass. Additionally, it uses default embeddings for the MSA and pairwise representations.

**Table A2:** Pearson correlations of lDDT, pLDDT and rASA.

| Method | Dataset | lDDT×rASA | pLDDT×rASA | lDDT×pLDDT |
| --- | --- | --- | --- | --- |
| **MSA** | **CASP13** | -0.276 | -0.313 | 0.786 |
|  | **CASP14** | -0.256 | -0.301 | 0.784 |
| **Template** | **CASP13** | -0.247 | -0.310 | 0.697 |
|  | **CASP14** | -0.257 | -0.300 | 0.686 |
| **Heuristic C-β** | **CASP13** | -0.198 | -0.211 | 0.779 |
|  | **CASP14** | -0.230 | -0.230 | 0.728 |
| **Template C-β** | **CASP13** | -0.302 | -0.318 | 0.656 |
|  | **CASP14** | -0.300 | -0.303 | 0.649 |
| **C36 (AF)** | **CASP13** | -0.182 | -0.179 | 0.681 |
|  | **CASP14** | -0.204 | -0.203 | 0.633 |
| **FASPR (AF)** | **CASP13** | -0.209 | -0.199 | 0.811 |
|  | **CASP14** | -0.242 | -0.225 | 0.758 |
| **AttnPacker** | **CASP13** | -0.265 | -0.155 | 0.341 |
|  | **CASP14** | -0.291 | -0.154 | 0.388 |
| **AttnPacker (AF)** | **CASP13** | -0.285 | -0.311 | 0.644 |
|  | **CASP14** | -0.298 | -0.301 | 0.637 |

Pearson correlations between lDDT, pLDDT and rASA for CASP13 and CASP14. AlphaFold2 refinement with: a full MSA and no template (*MSA*), full ground truth template (*Template*), ground truth backbone and the C-β placed with a heuristic (*Heuristic C- $\beta$* ), backbone and C-β from ground truth template (*Template C- $\beta$* ) or ground truth backbone and side-chains placed with CHARMM 36 force field (*C36 (AF)*), FASPR or (*FASPR (AF)*). Additionally, AttnPacker was evaluated before (*AttnPacker*) and after (*AttnPacker (AF)*) AlphaFold refinement.

**Table A3:**
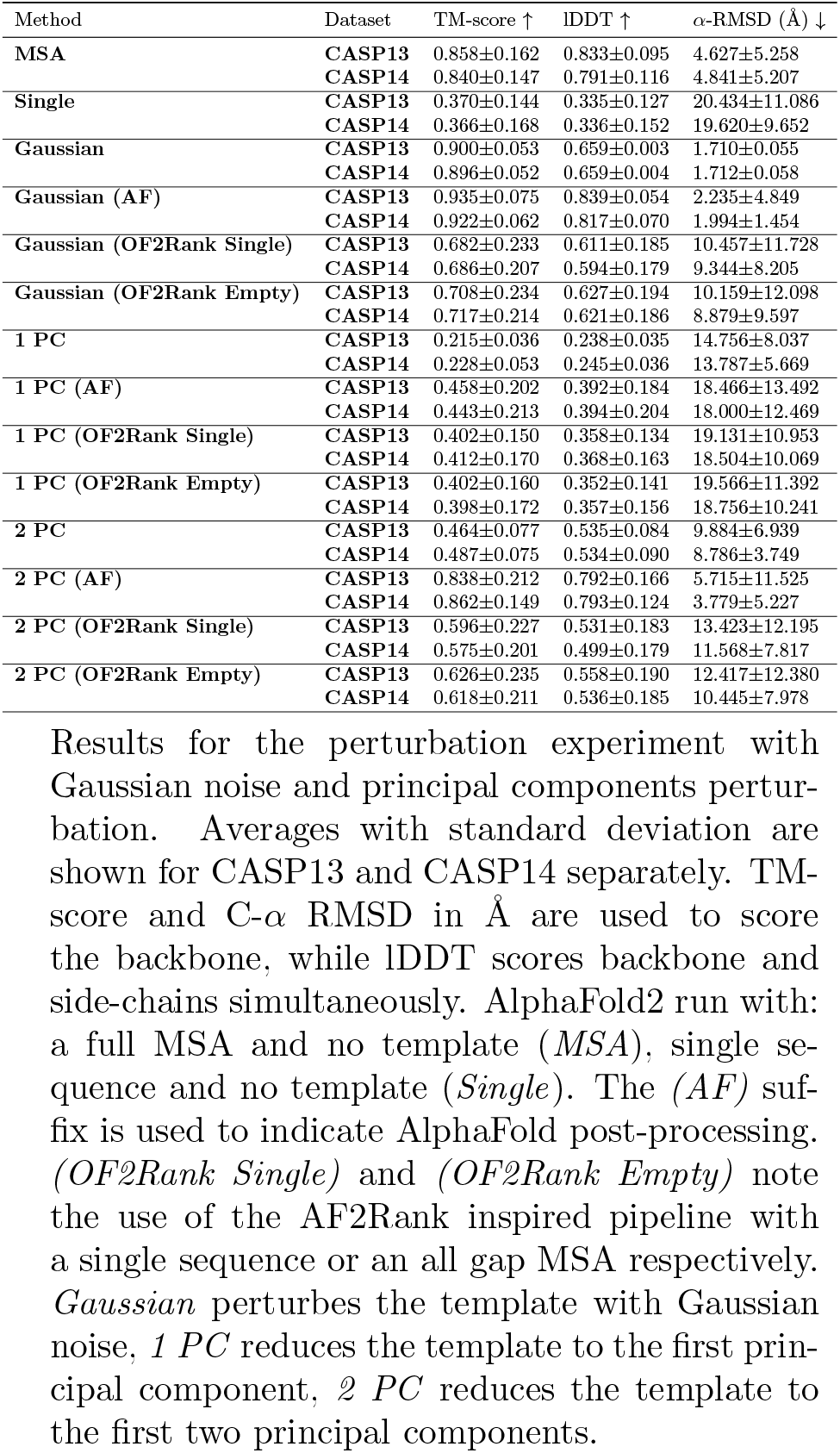
Refinement results on Gaussian noise and principal components perturbation.

**Table A4:**
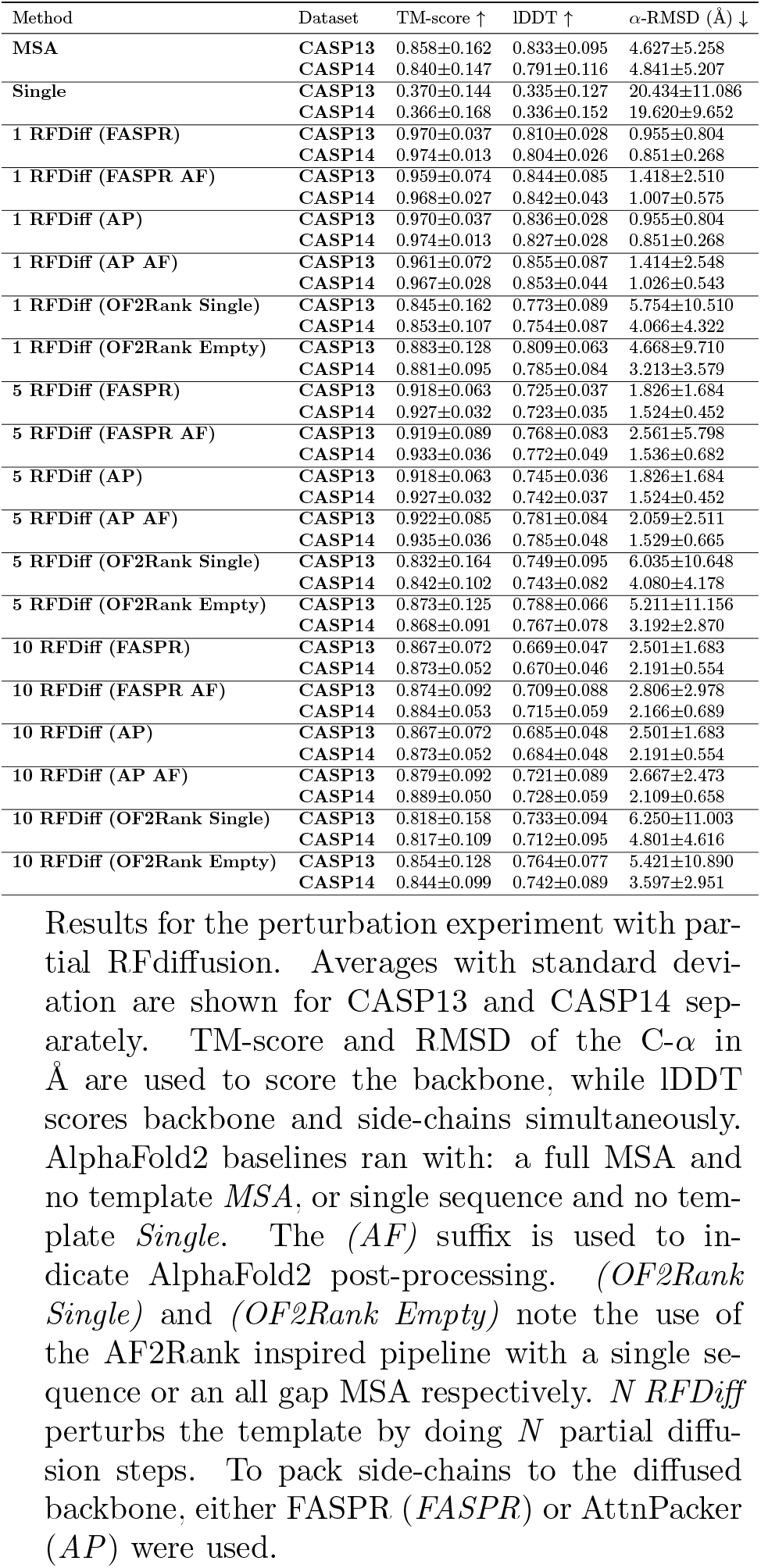
Refinement results on RFdiffusion perturbation.

**Table A5:** Refinement results with prev x modifications.

| Method | Dataset | TM-score ↑ | lDDT ↑ | α-RMSD ($\text{\AA}$) ↓ |
| --- | --- | --- | --- | --- |
| **MSA** | **CASP13** | 0.848±0.172 | 0.835±0.105 | 4.729±5.573 |
| | **CASP14** | 0.843±0.149 | 0.797±0.116 | 4.859±5.627 |
| **Single** | **CASP13** | 0.379±0.153 | 0.347±0.137 | 20.169±12.025 |
| | **CASP14** | 0.386±0.183 | 0.356±0.167 | 18.898±10.186 |
| **OF *prev_x*** | **CASP13** | 0.428±0.164 | 0.379±0.145 | 18.949±12.108 |
| | **CASP14** | 0.430±0.197 | 0.385±0.187 | 16.953±9.927 |
| **OF no *prev_x*** | **CASP13** | 0.853±0.180 | 0.832±0.113 | 4.914±6.223 |
| | **CASP14** | 0.848±0.146 | 0.794±0.116 | 4.732±5.885 |

Results for the *prev\_X* experiments. Averages with standard deviation are shown for CASP13 and CASP14 separately. TM-score and RMSD of the C- $\alpha$ in $\text{\AA}$ are used to score the backbone, while lDDT scores backbone and side-chains simultaneously. AlphaFold2 baselines ran with: a full MSA and no template *MSA*, or single sequence and no template *Single*. *OF prev\_x* indicates the results of an modified OpenFold version, where the ground truth template has been given as input for recycle 0 and the single sequence. An OpenFold version, where the *prev\_x* input is completely disabled and the standard amount of three iterations is run on a full MSA is shown in *OF no prev\_x*.

## Footnotes

1 https://github.com/ibmm-unibe-ch/template-analysis

2 https://github.com/aqlaboratory/openfold/pull/408

3 https://github.com/YoshitakaMo/localcolabfold

